## Supplementary Information for "Chromosomal genome assembly resolves drug resistance loci in the parasitic nematode *Teladorsagia circumcincta*"

Note on monepantel and genes associated with resistance	2

Supplementary Figure 1. Farm mean faecal egg count reduction test results, adjusted by L3 species proportions.	3

Supplementary Figure 2. Comparison of genetic variation between technical replicates.	4

Supplementary Figure 3. Comparison of chromosomal scaffolds between tci2_wsi3.0 and DNAZOO genome assemblies.	5

Supplementary Figure 4. Genetic differentiation between all samples.	6

Supplementary Figure 5. Relative sequencing coverage ratio along chromosome 5 between Farm 1 post:pre, Farm 2A post:pre and between Farm 1 post:Farm 2A post.	7

Supplementary Figure 6. Gene, BUSCO, and repeat density.	8

Supplementary Figure 7. Genome-wide gene coverage between susceptible and resistant isolates.	9

#### Note on monepantel and genes associated with resistance

None of the samples analysed were phenotypically resistant to monepantel. Monepantel was first registered for use in New Zealand in 2009 and then in Australia and the United Kingdom in 2010, and since then, resistance has evolved rapidly (David J. Bartley et al. 2019). Resistance has been proposed to be associated with loss-of-function variants in three genes – *mptl-1*, *deg-3* and *des-2* (Rufener et al. 2009) – with genetic variants found frequently in *mptl-1* of resistant *H. contortus* (Bagnall et al. 2017) and *T. circumcincta* (Turnbull et al. 2019) and, more recently, a genome-wide approach identified strong evidence of selection around a discrete region of the *H. contortus* genome containing the three genes (Niciura et al. 2019). In tci2_wsi3.0, we find *mptl-1* (TCIR_10044060) and *des-2* (TCIR_10044070) on chromosome 2 at ~15.1 Mb; however, no evidence of genetic differentiation is present in these data in that region. We could not find *deg-3* in our genome or annotation, which should lie between *mptl-1* and *des-1*. Given that orthologs of *deg-3* are found in the WASHU genome assembly (we detect two genes, TELCIR_11817 and TELCIR_18854, one of which may not be a complete gene), we conservatively conclude that the absence is due to an assembly artefact.

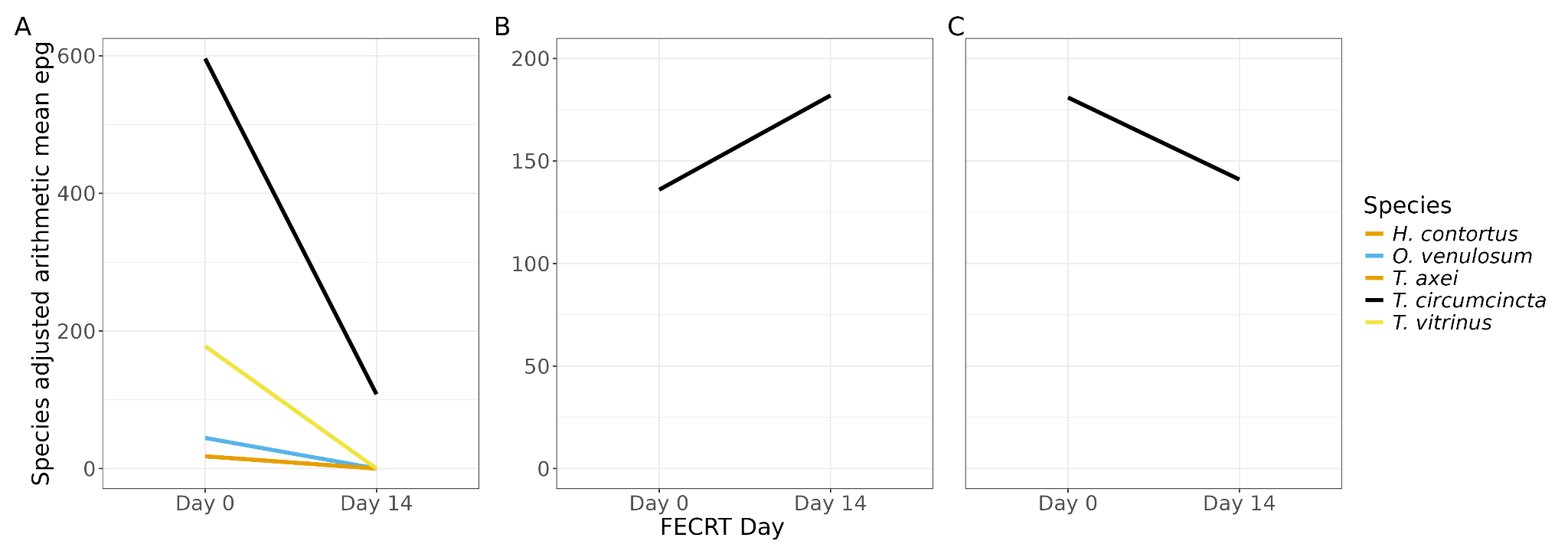

#### Supplementary Figure 1. Farm mean faecal egg count reduction test results, adjusted by L3 species proportions.

Data shown is the arithmetic mean of the faecal egg count reduction pre-to-post treatment, adjusted by L3 Trichostrongylid species proportions following PCR of the ITS2 region. A. Farm 1 ivermectin FECRT. B. Farm 2 benzimidazole FECRT. C. Farm 2 ivermectin FECRT. Species = *Haemonchus contortus*, *Oesophagostomum venulosum*, *Trichostrongylus axei, Teladorsagia circumcincta, Trichostrongylus vitrinus.* Key: epg = eggs per gram, FECRT = Faecal egg count reduction test

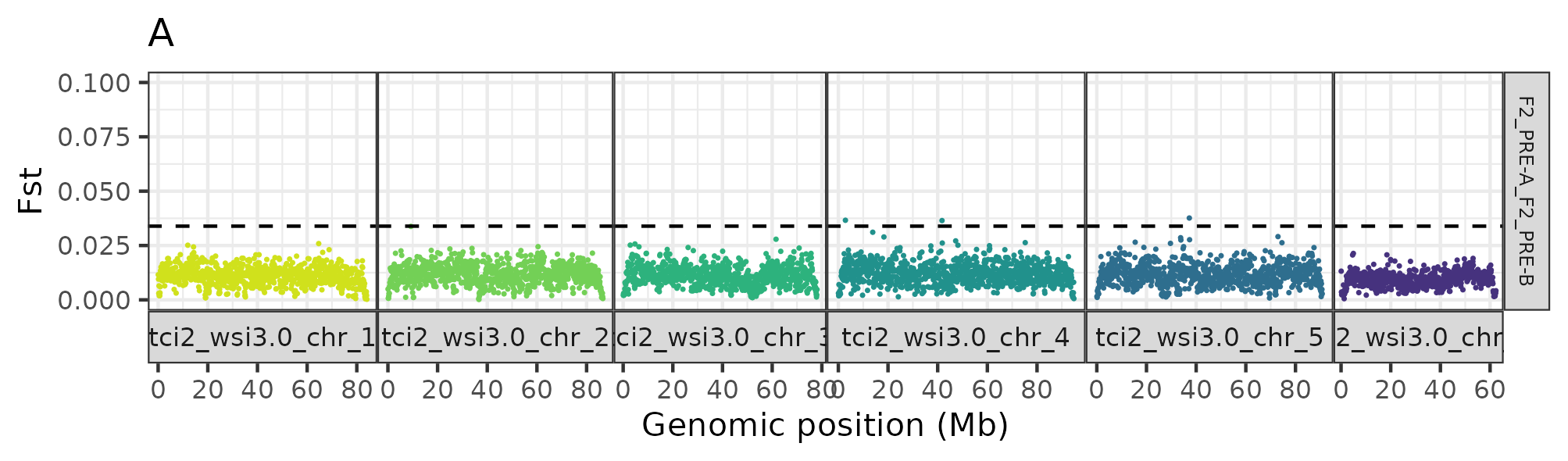

#### Supplementary Figure 2. Comparison of genetic variation between technical replicates.

DNA from Farm 2 samples was split into two sub-samples (“A” and “B” samples), from which individual libraries were generated and sequencing performed. Hence, comparing these samples provides a means to quantify the technical variation that drives differences between pairwise comparisons. Here, we have used this comparison to set a genome-wide level of significance, defined as the genome-wide mean *F*st + 5 standard deviations (SD) (GWsig = 0.03390286), which is used in Figure 2A to identify peaks of differentiation between pre- and post-treatment comparisons.

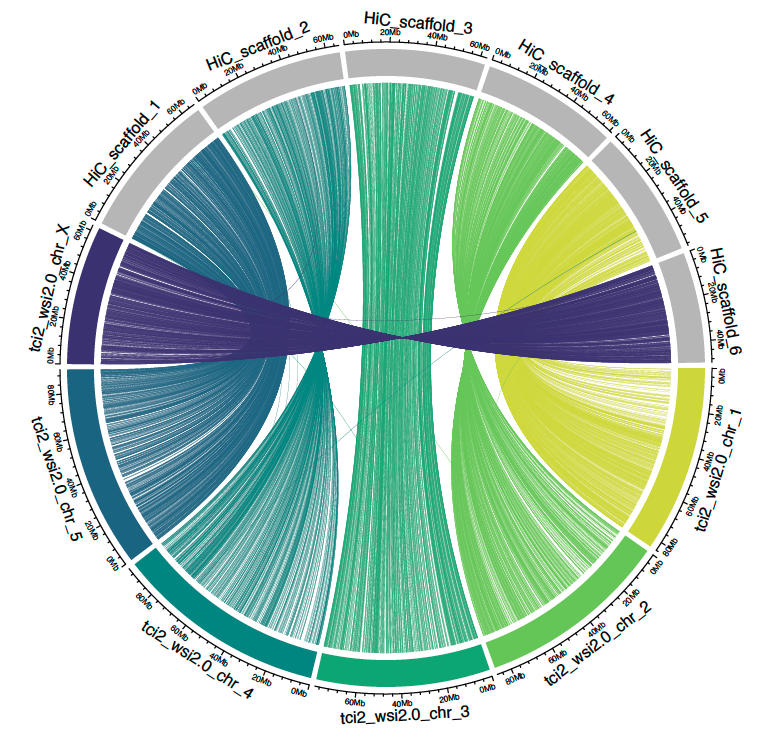

#### Supplementary Figure 3. Comparison of chromosomal scaffolds between tci2_wsi3.0 and DNAZOO genome assemblies.

The CIRCOS plot comparing the two genome assemblies demonstrates that, despite being derived from different strains and assembled independently, the chromosome structure of each genome assembly is consistent. The outer circle is divided into chromosomal segments proportionately sized and coloured by the tci2_wsi3.0 chromosome ID, whereas the DNAZOO scaffold segments are grey. The line segment shows positional information of shared nucleotide sequences connecting the two assemblies.

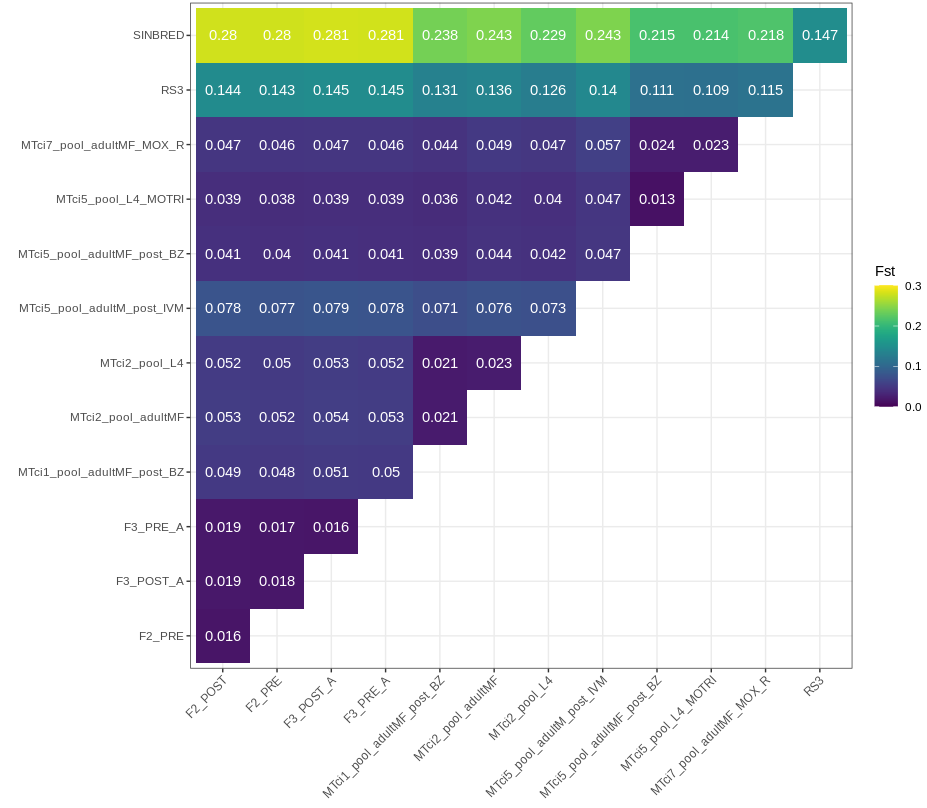

#### Supplementary Figure 4. Genetic differentiation between all samples.

Pairwise genetic differentiation (*F*st) is shown for all pairs of samples used to investigate anthelmintic resistance associated variation. These data highlight the differences between the NZ samples from Choi et al (Sinbred and RS3) from all other isolates which were obtained from the UK.

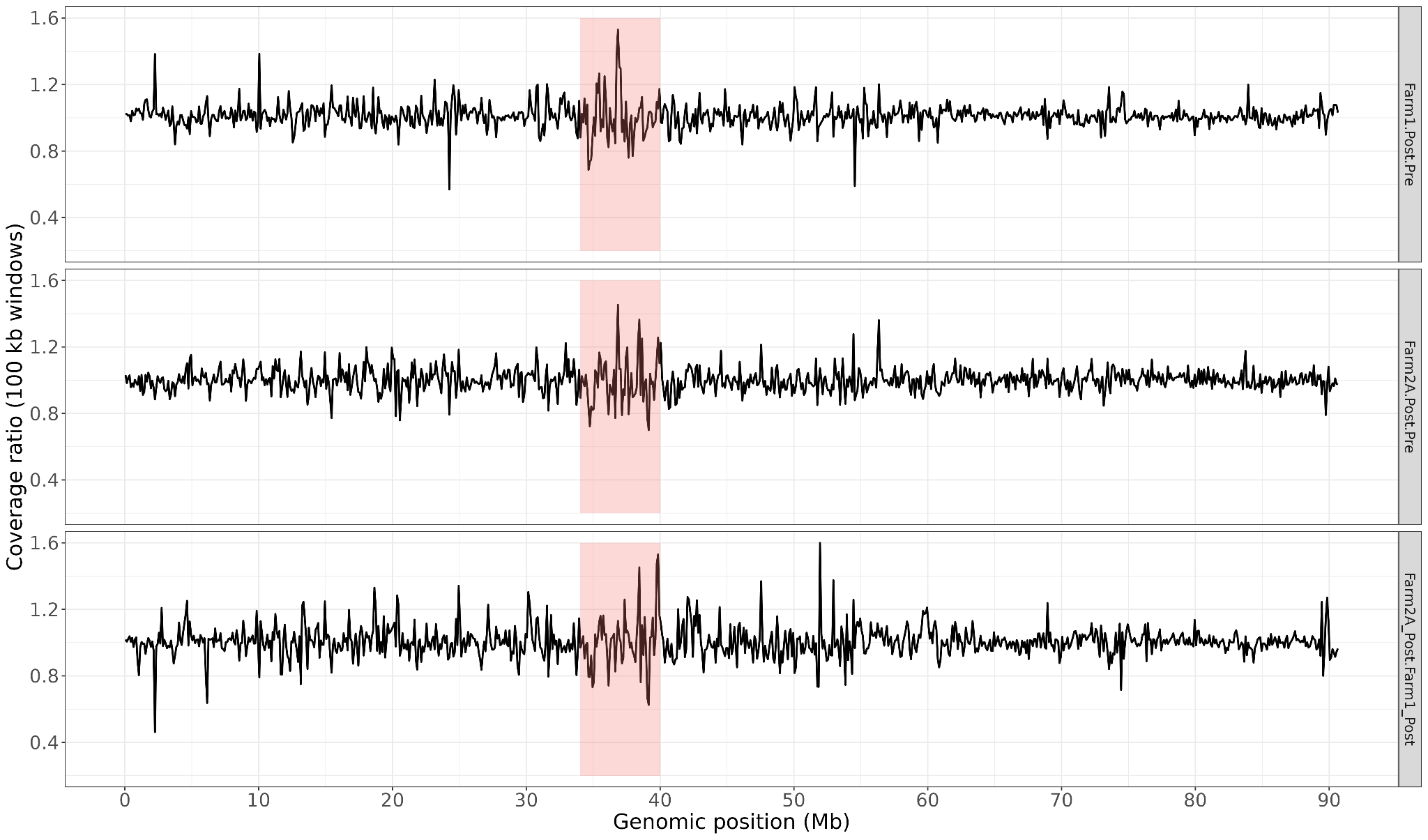

#### Supplementary Figure 5. Relative sequencing coverage ratio along chromosome 5 between Farm 1 post:pre, Farm 2A post:pre and between Farm 1 post:Farm 2A post.

The red rectangle delineates the ivermectin locus. Relative coverage ratio was determined after first dividing the 100 kb window average coverage by the chromosome 5 median depth coverage for that sample.

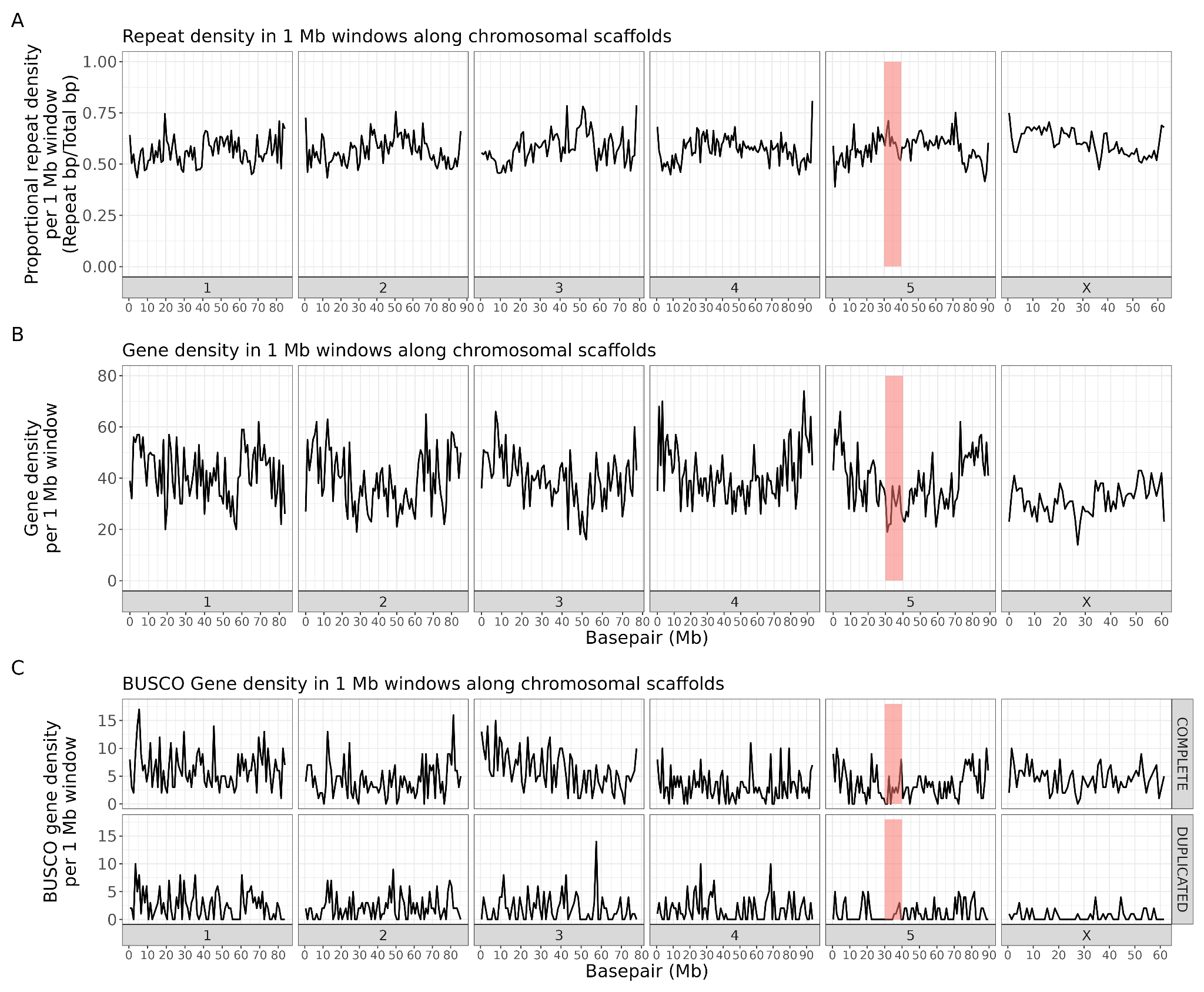

#### Supplementary Figure 6. Gene, BUSCO, and repeat density.

Repeat density (A), gene density (B) and density of ‘Complete’ or ‘Duplicated’ BUSCOs (C) from *compleasm* genome output were calculated in 1 Mb windows along each chromosome. For repeat density, the repeat proportion in the final window of each chromosome (less than 1 Mb) was adjusted by the window length, while for gene and BUSCO data, the final window data was removed. The red rectangle delineates the ivermectin locus.

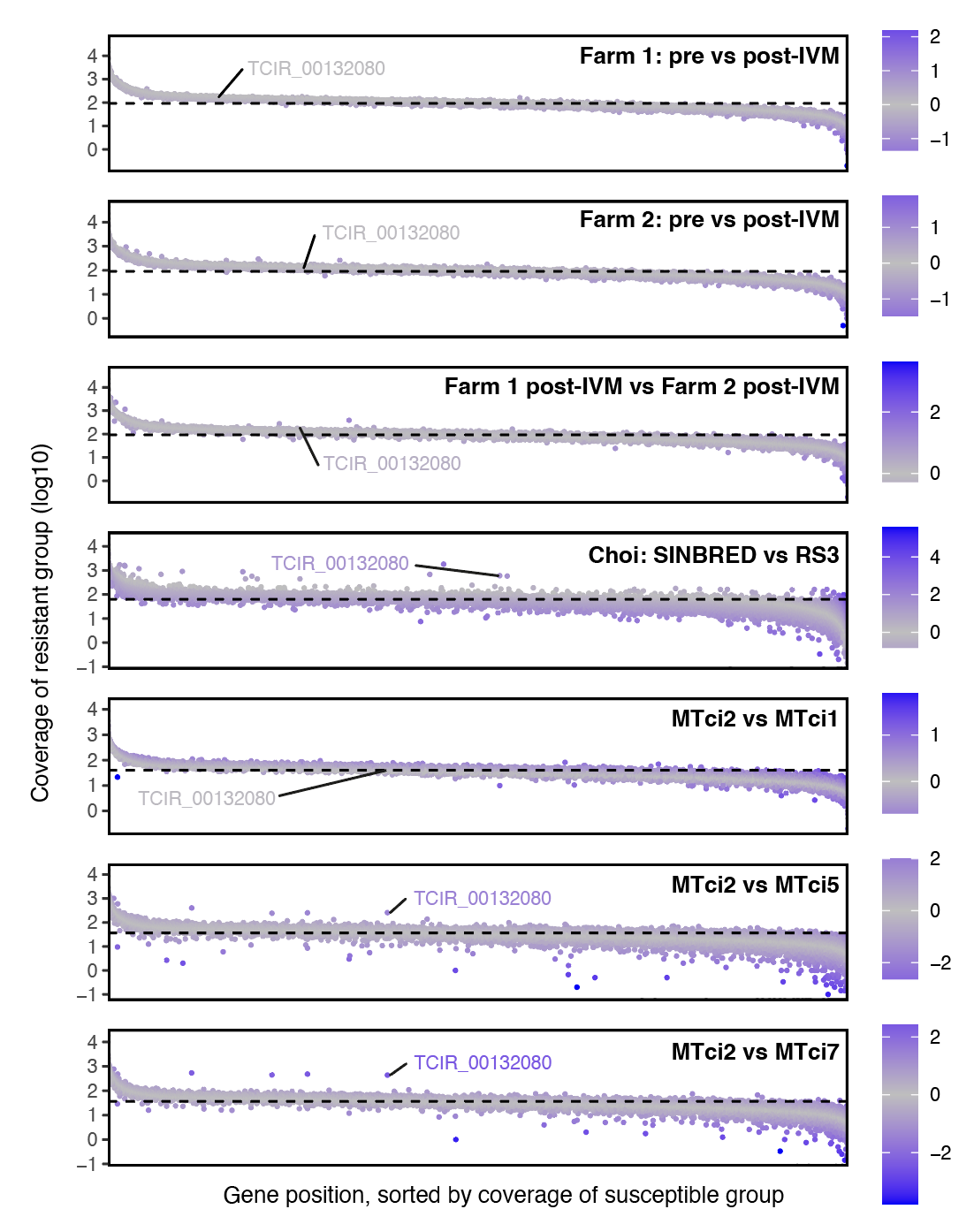

#### Supplementary Figure 7. Genome-wide gene coverage between susceptible and resistant isolates.

Each plot represents the distribution of sequencing read coverage of a resistant isolate relative to the expected coverage based on the susceptible isolate. For each comparison, coverage per gene is determined (mapped sequence reads per exon, from which a median coverage for all exons is calculated) for both susceptible and resistant isolates. Then, the order of the genes is sorted by coverage of the susceptible isolate from high-to-low coverage, after which the coverage of the resistant isolate is plotted (points, log10 coverage). This approach accounts for variation in sequencing coverage to reveal outliers in which coverage in the resistant isolates is higher or lower than expected. The points are coloured by the ratio of coverage (resistant/susceptible) to highlight outliers. In each plot, *pgp-9* (TCIR_00132080) is indicated, as it was the focus of the copy number variation analyses.
